# Quantifying full-length circular RNAs in cancer

**DOI:** 10.1101/2021.02.04.429722

**Authors:** Ken Hung-On Yu, Christina Huan Shi, Bo Wang, Savio Ho-Chit Chow, Grace Tin-Yun Chung, Ke-En Tan, Yat-Yuen Lim, Anna Chi-Man Tsang, Kwok-Wai Lo, Kevin Y. Yip

**Affiliations:** Department of Anatomical and Cellular Pathology, The Chinese University of Hong Kong, Shatin, New Territories, Hong Kong; Department of Computer Science and Engineering, The Chinese University of Hong Kong, Shatin, New Territories, Hong Kong; School of Life Sciences, The Chinese University of Hong Kong, Shatin, New Territories, Hong Kong; Institute of Biological Sciences, Faculty of Science, University of Malaya, Kuala Lumpur 50603, Malaysia; Hong Kong Bioinformatics Centre, The Chinese University of Hong Kong, Shatin, New Territories, Hong Kong; Hong Kong Institute of Diabetes and Obesity, The Chinese University of Hong Kong, Shatin, New Territories, Hong Kong

## Abstract

Circular RNAs (circRNAs) are abundantly expressed in cancer. Their resistance to exonucleases enables them to have potentially stable interactions with different types of biomolecules. Alternative splicing can create different circRNA isoforms that have different sequences and unequal interaction potentials. The study of circRNA function thus requires knowledge of complete circRNA sequences. Here we describe psirc, a method that can identify full-length circRNA isoforms and quantify their expression levels from RNA sequencing data. We confirm the effectiveness and computational efficiency of psirc using both simulated and actual experimental data. Applying psirc on transcriptome profiles from nasopharyngeal carcinoma and normal nasopharynx samples, we discover and validate circRNA isoforms differentially expressed between the two groups. Compared to the assumed circular isoforms derived from linear transcript annotations, some of the alternatively spliced circular isoforms have 100 times higher expression and contain substantially fewer microRNA response elements, demonstrating the importance of quantifying full-length circRNA isoforms.

## 1 Introduction

CircRNAs are a class of single-stranded RNAs with the 5’ and 3’ ends covalently linked [1–3]. Although they have been discovered for a long time [4, 5], research in circRNAs has only been reinvigorated in recent years by the discoveries that some circRNAs are highly abundant [6, 7] and conserved across species [8], and have regulatory potentials by functioning as microRNA (miRNA) sponges [9, 10]. Several additional sequence-or structure-specific functions of circRNAs have also been proposed [1, 3, 11–13].

A number of circRNAs are highly expressed in cancer or are differentially expressed between cancer and normal tissues [14]. Some of them have been shown to play oncogenic [15] or tumor-suppressive [16] roles. Since circRNAs are relatively stable as compared to their linear counterparts due to their resistance to RNA exonuclease [10], they can potentially be used as diagnostic biomarkers of cancer [17, 18].

Currently, the standard way of detecting circRNAs genome-wide is RNA sequencing (RNA-seq). Since circRNAs are not polyadenlynated, protocols without poly-A enrichment are used, such as those that involve ribosomal RNA (rRNA) depletion. The resulting data contain a mixture of sequencing reads from both linear and circular transcripts. A usual way to enrich for circRNA reads is to apply RNase R treatment, which preferentially digests linear transcripts [10, 19].

Regardless of the RNA-seq protocol, a common step in identifying circRNAs from the sequencing data is to look for back-splicing junctions (BSJs), i.e., junctions that connect the 3’ end of a downstream exon to the 5’ end of an upstream exon, which indicate potential circularization events. Several computational methods have been proposed for identifying circRNAs from RNA-seq data using this idea [10, 20–36]. Major differences among these methods include their read alignment strategy, signals used to detect BSJs, dependency on annotations of linear transcript isoforms, and ability to distinguish circRNAs from other events that may also create unexpected junctions, such as lariats, fusion genes and tandem duplications. These methods have been extensively compared using benchmark data sets [37–39]. The identified circRNAs and their expression information are cataloged in several databases [18, 34, 40–44].

Similar to linear transcripts, circular transcripts can also be alternatively spliced to create different isoforms with the same BSJ [34, 45]. Four major types of alternative splice site selection, namely cassette exon, intron retention, alternative 5’ splice site and alternative 3’ splice site, can all be found in circRNAs [34]. Since the function of a circRNA depends strongly on its exact sequence, for example the presence of miRNA response elements (MREs) and binding sites of RNA-binding proteins, it is critical to resolve full-length circRNA sequences and quantify their expression levels [46].

Most existing circRNA detection methods can only identify BSJs but cannot determine full-length circular transcripts. The only existing methods that can identify and quantify fulllength circRNA transcripts are Sailfish-cir [26] and CIRI-full [36]. Sailfish-cir is not a complete pipeline for circRNA detection but rather depends on the back-splicing junctions produced by another method as input. CIRI-full’s full-length transcript identification method detects mostly short transcripts such as those shorter than the total length of the two sequencing reads in a mate pair [36], while longer transcripts could be missed.

In this paper, we propose psirc (<u>ps</u>eudo-alignment identification of c<u>irc</u>ular RNAs), the first complete pipeline that can detect full-length circRNA transcript isoforms of all lengths and quantify their expression levels directly from RNA-seq data. The main ideas are to use pseudo-alignment to efficiently identify potential BSJs and full-length isoforms, and a transcript de Bruijn graph (T-DBG) [47] to quantify both full-length linear and circular transcripts at the same time by means of likelihood maximization. Using simulations and RNA-seq, RT-PCR and quantitative RT-PCR (RT-qPCR) data from human cell line and tissue samples, we show that psirc can accurately identify and quantify BSJs and full-length transcript isoforms. We also use psirc to identify circRNAs differentially expressed between nasopharyngeal carcinoma (NPC) and normal nasopharynx (NP) samples with independent experimental validations. Finally, we find that some of the alternatively spliced circular isoforms have much higher expression levels and fewer MREs than their corresponding “default” isoforms based on linear transcript annotations, which demonstrates that functional studies could be seriously hampered if these default isoforms are assumed without identifying and quantifying the actual full-length circular isoforms.

## 2 Results

### 2.1 The psirc method

The overall procedure of psirc consists of three steps, namely 1) identifying BSJs, 2) determining potential full-length transcript isoforms, and 3) quantifying the expression levels of full-length transcript isoforms (Figure 1, Materials and Methods).

**Figure 1:**
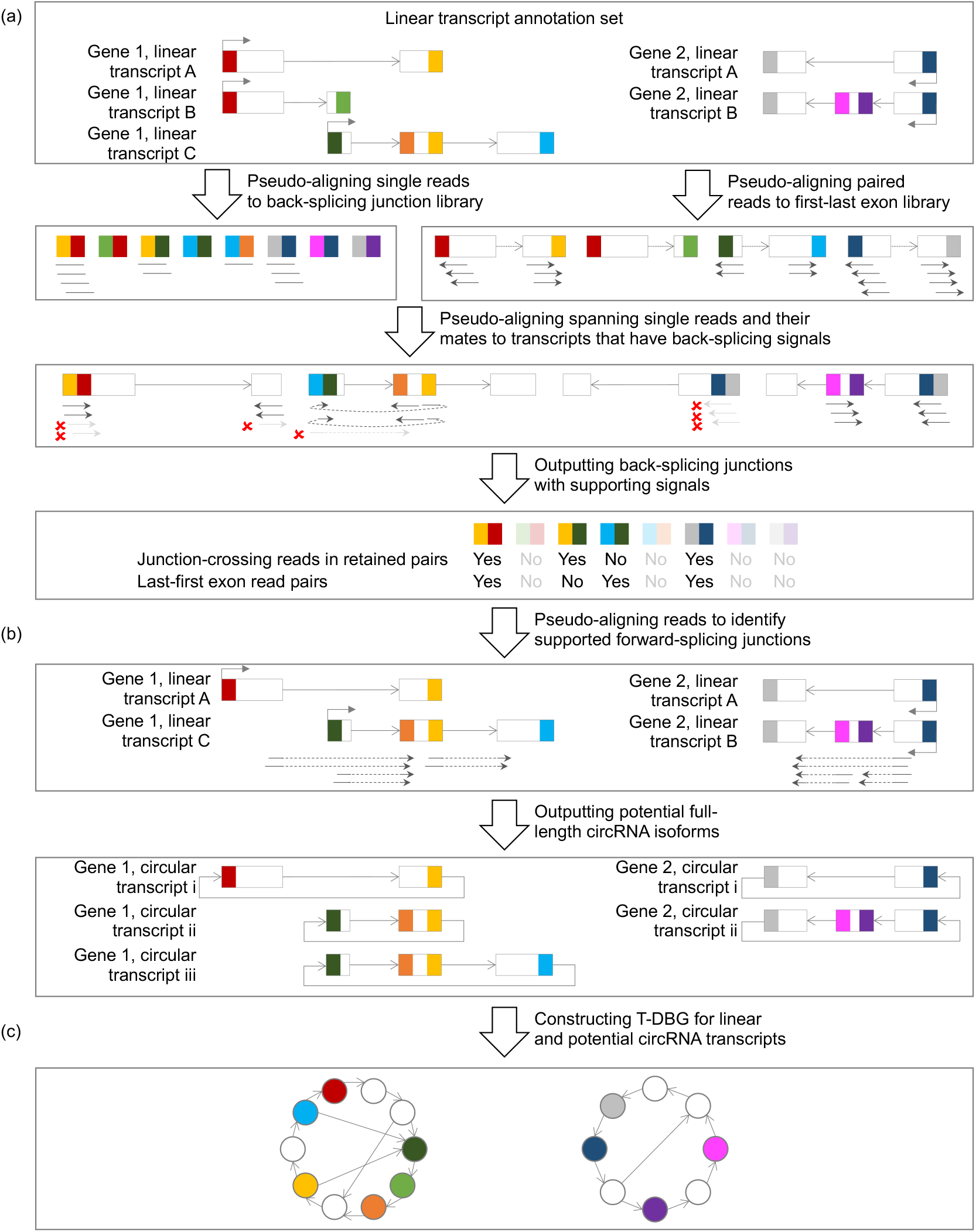
The psirc method. In the first step (a), sequencing reads are pseudo-aligned to potential BSJs in single-end mode and to the first and last exons of each transcript in paired-end mode. For each read that is pseudo-aligned to a BSJ (by means of rotating the end of the downstream exon to the beginning of the upstream exon), if its mate read is not pseudo-aligned to the same transcript or not pseudo-aligned in the opposite orientation, both of them will not be considered to support the BSJs (arrows in light gray with a red cross next to them). In the second step (b), the potential circRNA transcript isoforms are determined as those with all forward-splicing and backward-splicing junctions supported by sequencing reads. In the third step (c), a T-DBG is constructed for all linear and circular transcript isoforms for estimating their expression levels by another round of pseudo-alignment.

In the first step (Figure 1a), two types of signals are used to identify BSJs, namely junction-crossing reads and last-first exon read pairs. A junction-crossing read is a sequencing read split-pseudo-aligned to cover the 3’ most positions of a downstream exon and the 5’ most positions of an upstream exon. A last-first exon read pair is a read pair respectively pseudo-aligned to the first and last exons of an annotated linear transcript in an outward-facing manner. Based on the candidate BSJs identified by these signals, the transcript sequences involved are rotated for another round of pseudo-alignment in which the previous junction-crossing reads are expected to pseudo-align to them directly without split-alignments. The candidate BSJs confirmed by such pseudo-alignments are outputted to a file as an intermediate product, which can be used for analyses that do not require full-length circRNA isoforms.

In the second step (Figure 1b), all RNA-seq reads are pseudo-aligned to the linear transcripts that contain both the defining exons of any BSJ. The potential full-length circRNA transcript isoforms are then generated as those with all the involved forward- and backward-splicing junctions supported by sequencing reads using a graph searching algorithm. Since the BSJs identified in the first step of psirc can be defined by any pairs of exons, psirc allows the detection of circRNA isoforms that contain exon combinations not identical to any annotated linear isoforms.

Finally, in the third step (Figure 1c), a T-DBG is constructed for all linear and potential circular transcript isoforms. The pseudo-alignments of sequencing reads are then used to quantify the expression level of each linear and circular transcript isoform by likelihood maximization, with boundary effects taken care of by adjusting effective transcript lengths (Materials and Methods). To ensure only relevant sequencing reads are used to quantify the expression level of a transcript, we additionally i) check the orientations and insert sizes of the pseudo-alignments of read pairs, which for instance allow us to handle the reverse-overlap (RO) cases [36], ii) verify that reads that span BSJs are properly aligned on both sides, rather than assuming it based on the positional information of pseudo-alignments alone, and iii) consider the number of repeated sequences within a transcript that may lead to non-unique pseudo-alignments.

In all three steps, full alignments of sequencing reads are avoided by using kallisto [47] to perform pseudo-alignments, which makes psirc highly efficient in terms of both running time and memory consumption.

### 2.2 Effective BSJ detection with low time and memory requirements

We first verified the ability of psirc in identifying BSJs using three data sets. The first data set contained rRNA-depleted RNA-seq data from 11 human fetal samples (Table S1), from which circRNAs were identified previously [29]. We used psirc and three other methods to identify BSJs from each sample. These methods were 1) CIRI2 [23] and 2) CIRCexplorer2 [34], two methods that consistently ranked top in benchmarking studies [37–39], and 3) CircMarker [27], a method aiming at achieving high speed efficiency by using k-mer matching to avoid expensive sequence alignments.

From the results (Figure S1), CircMarker, CIRI2 and psirc identified comparable numbers of BSJs, while CIRCexplorer2 identified substantially less. For the former three methods, 80% or more of their identified BSJs were also identified by at least one other method. This high consistency among the three methods suggests that at least some of the BSJs identified by them but not CIRCexplorer2 were legitimate, and thus they achieved higher sensitivity than CIRCexplorer2 on this data set.

To evaluate the precision of the four methods, we tested them on two additional data sets, which contained matched rRNA-depleted and RNase R-treated RNA-seq data from human HeLa and Hs68 cells, respectively (Table S1). We applied each method to identify BSJs from the rRNA-depleted RNA-seq data, and then used the corresponding RNase R treated RNA-seq data to classify these junctions into Enriched, Unaffected, Depleted, and Abolished cases, defined as junctions with increased, similar, decreased but non-zero, and zero support in the RNase R data, respectively (Materials and Methods).

After combining the replicates, the numbers of BSJs identified by CircMarker, CIRI2 and psirc were again larger than those of CIRCexplorer2 (Figure S2a,b). We then computed precision values based on four different definitions, depending on whether Unaffected cases were considered true positives and whether Depleted cases were considered false positives, since these cases were more ambiguous in general. Overall, psirc had precision values comparable to CIRCexplorer2 and CIRI2 and higher than CircMarker (Figure S2c,d).

In terms of computational cost, when running with 25 threads, psirc required the least amount of resources among the four methods in terms of elapsed time, CPU time (summing over all threads) and memory (Figure S3). For every sample, compared to the second best method, psirc consistently required only one-third of its CPU time and memory. To test the applicability of psirc when computational resources are limited, we also ran it using only one thread, and found that the resulting elapsed time was still comparable to CircMarker and faster than CIRCExplorer2 when these methods were run with 25 threads, while the CPU time of psirc was further substantially reduced to one third as compared to running it with 25 threads (Figure S3).

Together, these results show that in terms of identifying BSJs, psirc requires much less computational resources but still achieves comparable sensitivity and precision as the best of the other three methods.

### 2.3 Accurate quantification of full-length circRNA isoforms

We next tested the ability of psirc in quantifying expression levels. First, we used the data set of human fetal samples mentioned above, which also included the expression levels of some BSJs independently measured by RT-qPCR [29]. We applied psirc to deduce the expression level of each full-length circRNA isoform using the RNA-seq data, based on which we computed the expression level of each BSJ by aggregating the expression levels of all the full-length circRNA isoforms that involved this junction. We then compared these deduced BSJ expression levels with those measured by RT-qPCR. For benchmarking purpose, we also deduced BSJ expression levels directly from the RNA-seq data using the other three methods.

From the results, for all four methods, the deduced BSJ expression levels were strongly correlated with the RT-qPCR results no matter the read counts were normalized (Figure S4a) or not (Figure S4b). Among the four methods, the correlation values were slightly stronger for psirc than the other three methods.

In the above comparisons, each method was evaluated based on the BSJs identified by it, which could be different from the BSJs identified by the other methods. To compare the quantification capability of the different methods on the same ground, we designed a novel benchmarking procedure. First, we divided the whole BSJ quantification process into three components, namely BSJ calling (*B*), full-length circRNA isoform inference (*F*), and expression level quantification (*Q*). The quantification component was further divided into two types, namely quantification of BSJs without inferring full-length isoforms (*Q*_*b*_) and quantification of BSJs by quantifying and aggregating full-length circRNA isoform expression levels (*Q*_*f*_). Some of the methods provided all three components while the others provided only some of them (Table S2). We then established full pipelines by mixing-and-matching components of different methods, such that the quantification performance of different methods could be fairly compared based on the same set of BSJs or full-length transcript isoforms. For this part of analysis, we also included Sailfish-cir since BSJs identified by other methods could be supplied to it as input. In addition, we included two new methods of the CIRI series, CIRI-full [36] and CIRI-quant [35], to provide supplementary or alternative components for CIRI2.

Using the benchmarking procedure, we first compared the quantification performance of four pipelines based on the BSJs identified by CIRI2 (Figure 2a-d). Among the three pipelines that computed BSJ expression levels by aggregating expression levels of full-length isoforms (i.e., those with the *Q*_*f*_ component), two of them achieved stronger correlations than the pipeline that computed BSJ expression levels directly from BSJs (i.e., the one with the *Q*_*b*_ component). The only exception was the pipeline involving CIRI-full, which was unable to infer any full-length transcript isoform for many BSJs since it was designed to identify short isoforms. Among the four pipelines, the one with expression quantification performed by psirc achieved the strongest correlation of -0.835.

**Figure 2:**
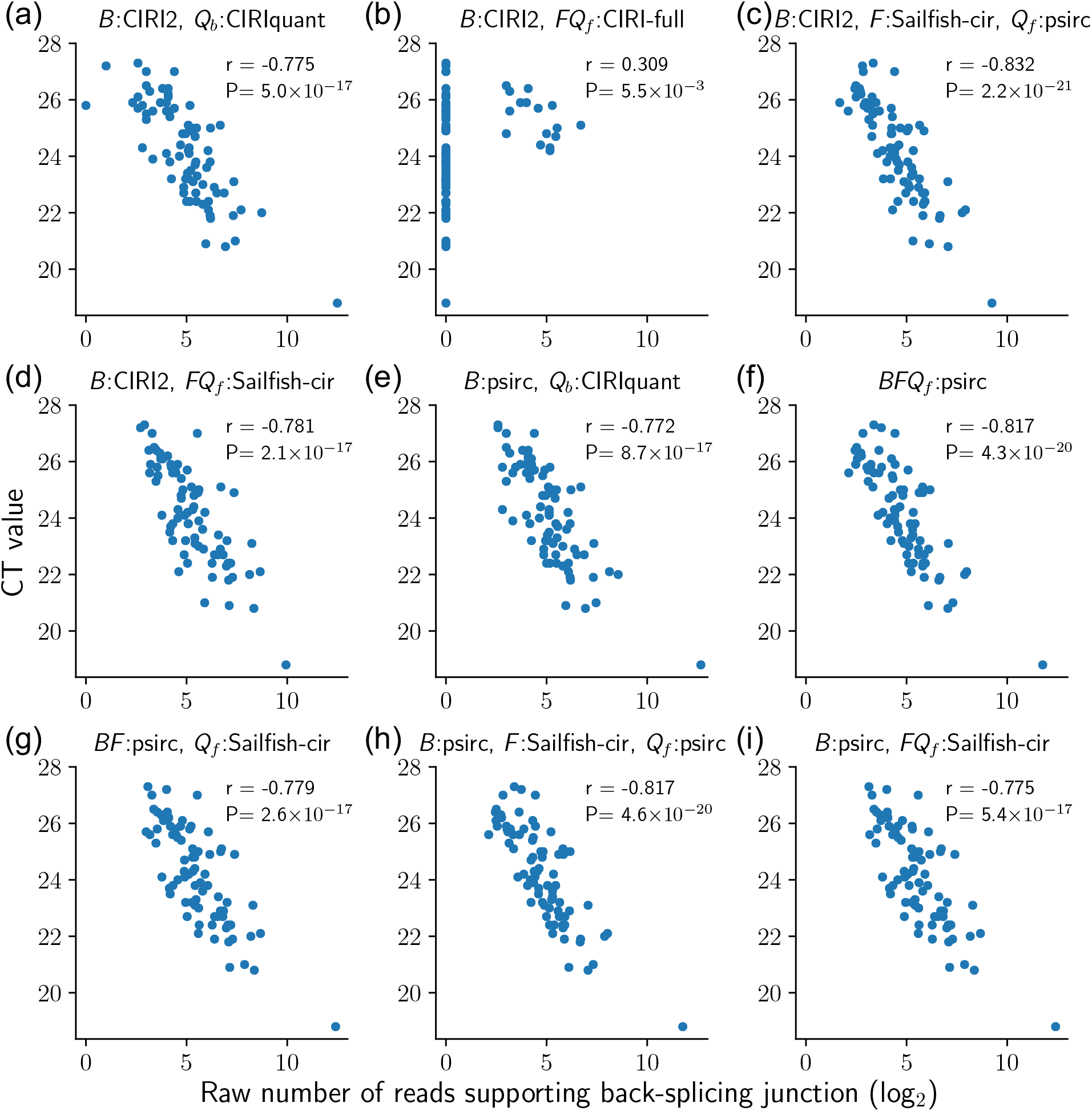
Comparison of different pipelines based on their determined read counts that support each BSJ from the human fetal samples. Each point corresponds to one BSJ in one sample. Each panel corresponds to a different pipeline by combining the three components from different methods. If the same method was used for multiple components, they are written together. For example, “*B*:psirc, *FQ*_*f*_ :Sailfish-cir” means the BSJs were identified by psirc, while both the full-length transcript isoforms and their expression levels were deduced by Sailfish-cir. All pipelines involving full-length quantification (*Q*_*f*_) aggregated the expression levels of all transcripts that involved a BSJ into the expression level of the junction.

Similarly, when the BSJs were identified by psirc (Figure 2e-i), the four pipelines involving the inference of full-length transcripts achieved stronger correlations than the pipeline that quantified the BSJs directly. Among these four pipelines, when the full-length isoform inference method was fixed, using psirc for quantification achieved stronger correlations than Sailfish-cir (−0.817 for “*BFQ*_*f*_ :psirc” versus -0.779 for “*BF* :psirc, *Q*_*f*_ :Sailfish-cir”; -0.817 for “*B*:psirc, *F* :Sailfish-cir, *Q*_*f*_ :psirc” versus -0.775 for “*B*:psirc, *FQ*_*f*_ :Sailfish-cir”).

All these conclusions still hold when the deduced BSJ expression levels were normalized (Figure S5). Overall, these results show that the full-length circRNA isoform inference and quantification of psirc enabled it to deduce BSJ expression levels that were strongly correlated with RT-qPCR results.

Next, we set forth to evaluate psirc’s accuracy in quantifying the expression levels of full-length circRNA isoforms. Since large-scale experimental data of full-length circRNA expression levels are not available, we performed this evaluation using two sets of simulated data. We benchmarked the results of psirc against those produced by Sailfish-cir on the basis of its good performance in quantifying BSJs in the evaluations above. To directly compare the quantification component of the two methods, we supplied the full-length transcript isoforms inferred by psirc as the common input to both methods.

In the first set of simulated data, we followed Li et al.’s original approach to testing Sailfish-cir [26] to simulate 11 groups of genes. Each group contained 500 genes with one linear isoform and one circular isoform having independent expression levels, leading to 1000 isoforms per group. The 11 groups differed by the degree of overlap between the linear and circular sequences, ranging from 0% to 100% (Figure S6). When the overlap ratio was low, both psirc and Sailfish-cir were able to estimate expression levels accurately, with the correlation between the estimated and actual expression levels of the 1000 isoforms in each group as high as 0.99 (Figure 3a,b). However, when the overlap ratio was 100%, the correlation value of Sailfish-cir dropped to below 0.93 while that of psirc remained higher than 0.99. To check whether psirc’s high correlation was due to biases caused by improper data normalization, we plotted transcript expression levels against their lengths for the group with 100% sequence overlap (Figure 3c) but did not observe any obvious correlation between expression level and gene length that could have caused biases. We further plotted the estimated and actual read counts of this group of isoforms (Figure 3d) and found that Sailfish-cir tended to over-estimate read counts of linear transcripts and under-estimate those of circular transcripts, which could not be shown in the previous BSJ quantification results that involved only circular transcripts. In contrast, psirc was able to quantify both linear and circular transcripts accurately.

**Figure 3:**
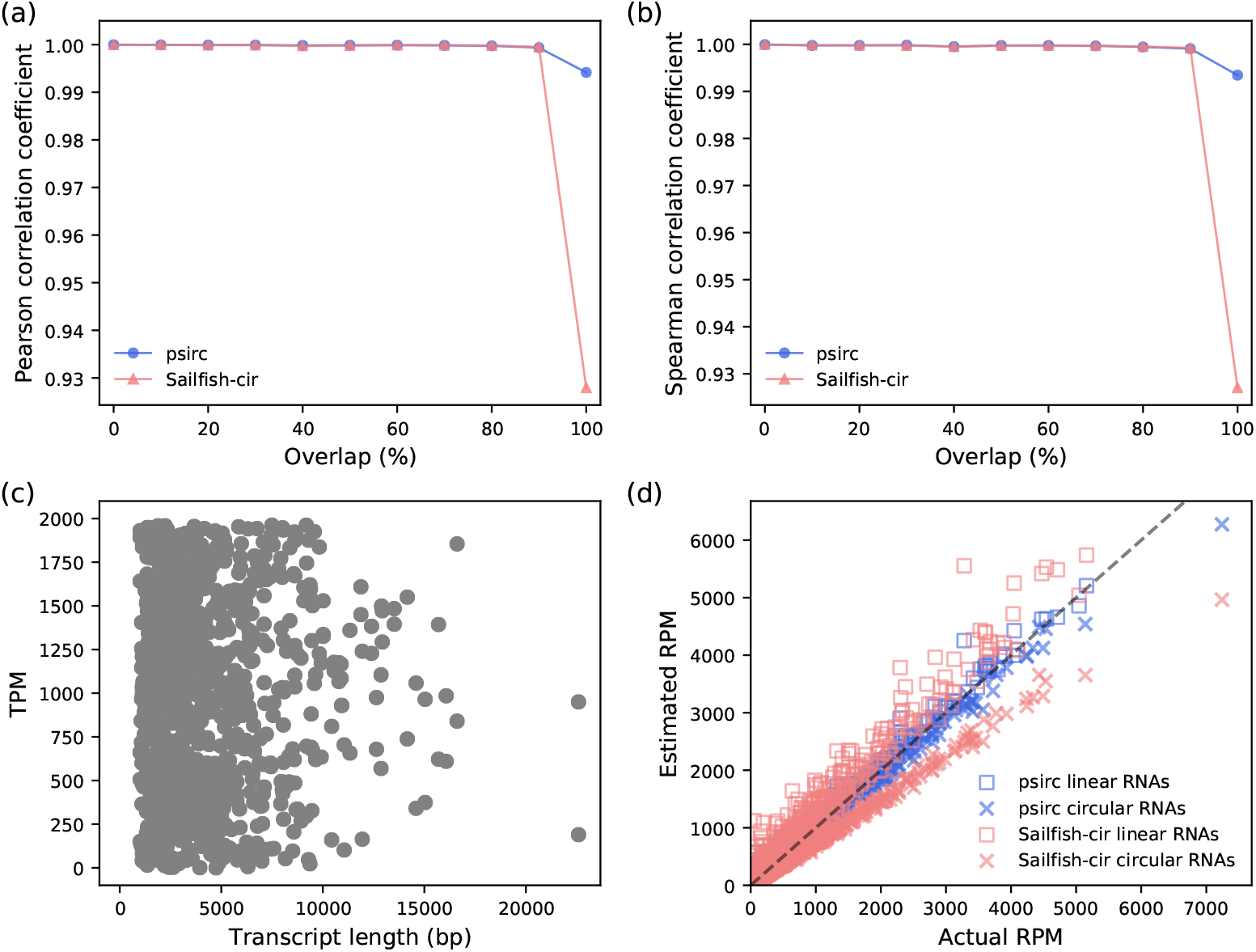
Quantification performance of psirc and Sailfish-cir on the first simulated data set. Pearson (a) and Spearman (b) correlation coefficients were computed between the estimated and actual expression levels for the 1000 transcript isoforms in each of the 11 groups. For the group with 100% sequence overlap between linear and circular transcript isoforms, the scatter plots show the actual expression levels in transcripts per million reads aligned (TPM) against transcript lengths (c), and the read counts per million reads aligned (RPM) estimated by the two methods against the actual read count of each transcript isoform (d).

To test whether psirc can handle more complex gene structures, we produced a second set of simulated data. In this set of data, there were 10 groups of genes. Each gene in the *i*-th group had *i* linear transcript isoforms and the same number of circular isoforms with exactly the same sequences as the linear counterparts (i.e., 100% sequence overlap) but independent expression levels. We produced data for 10 random genes in each group, and repeated the procedure 10 times. For all 10 groups, psirc outperformed Sailfish-cir by a clear margin (Figure 4a,b). In general, the performance of both methods dropped as the number of transcript isoforms per gene increased. Yet importantly, the correlation between psirc’s estimated expression levels and the actual expression levels remained higher than 0.9 even when there were 10 linear and 10 circular isoforms per gene. When we plotted the estimated and actual read counts of the isoforms (Figure 4c-i), again we observed that the quantification results of psirc were closer to the actual values than were the results of Sailfish-cir in all cases, even the genes chosen to be included in the plots were already the ones that Sailfishcir performed the best in each group.

**Figure 4:**
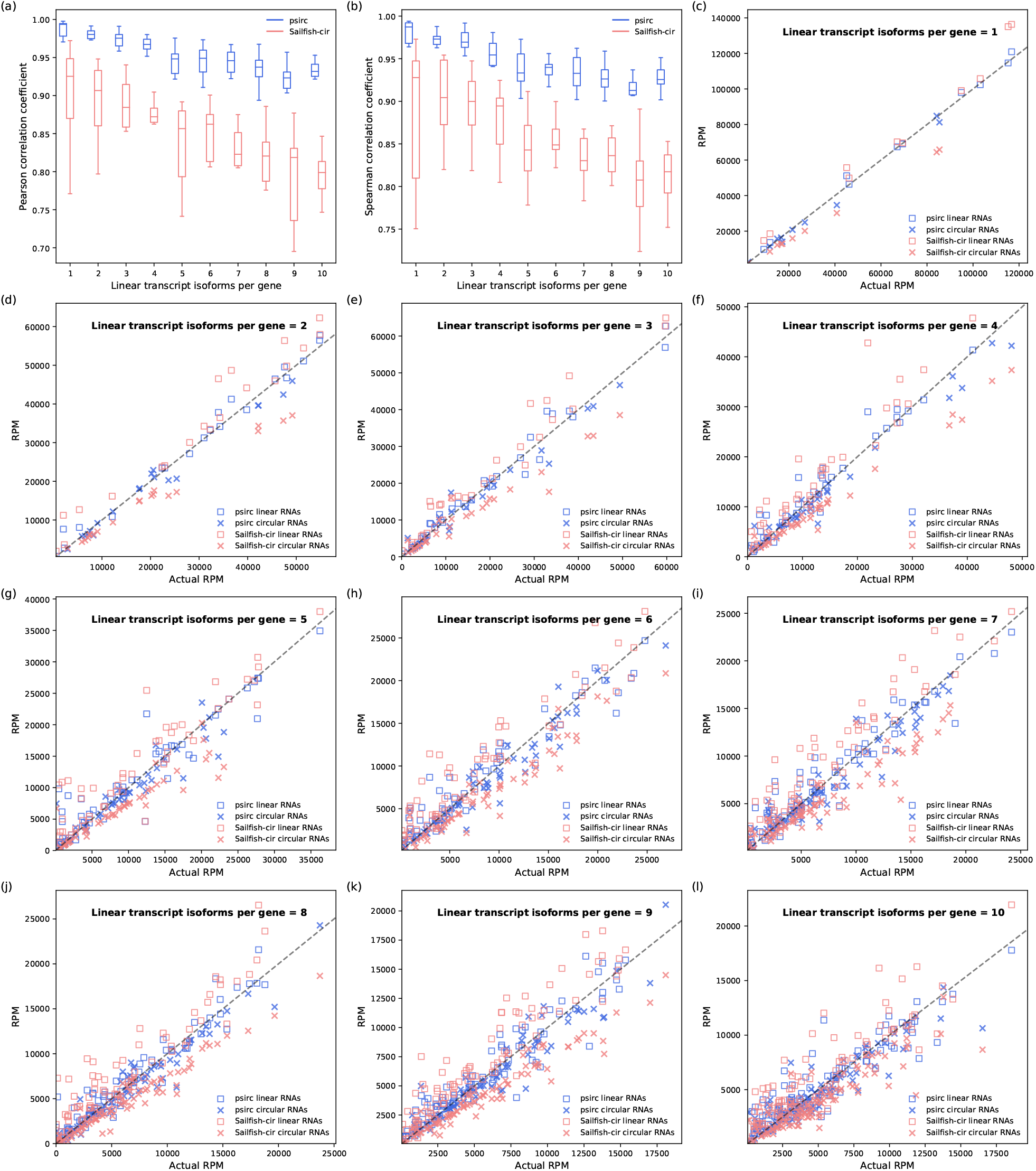
Quantification performance of psirc and Sailfish-cir on the second simulated data set. Pearson (a) and Spearman (b) correlation coefficients were computed between the estimated and actual expression levels. Each box plot shows the distribution of correlations from the 10 sets of random transcripts, with each correlation coefficient computed based on all the transcripts from the 10 random genes in that set. (c-i) The estimated and actual read counts per million reads aligned are shown for each transcript isoform for the gene that Sailfish-cir achieved the strongest Pearson correlation in each group.

### 2.4 Discovery of cancer-related circRNAs in nasopharyngeal carcinoma

With the effectiveness and efficiency of psirc verified by the series of tests above, we applied it to identify BSJs and full-length circRNA transcript isoforms and deduced their expression levels from rRNA depleted RNA-seq data of 11 NPC cell lines, xenografts and patient tumor specimens and 4 NP cell lines (Table S3). Sequencing reads were aligned to both the human and Epstein-Barr Virus (EBV) genomes at the same time, allowing for a joint detection and quantification of both human and EBV full-legnth circRNAs.

We detected 8,401-28,809 BSJs from the different samples, from which 8,590-29,645 full-length circRNA isoforms were inferred. Focusing on only the frequently expressed cases, defined as those expressed in at least 70% of NPC or NP samples, 3,145-5,862 BSJs and 2,024-3,965 full-length isoforms were identified from the samples. (Table 1, Supplementary Files 1-4)

**Table 1:** Numbers of human and EBV BSJs and full-length circular isoforms identified by psirc from the NPC and NP samples. The samples are listed in the same order as in Table S3, with the NPC samples listed before the NP samples. The total numbers identified (i.e., with read supports) from each sample are listed in the first two columns. The last two columns consider only BSJs and isoforms expressed in at least 70% of the NPC or NP samples.

| Sample | Identified |  | Among commonly expressed |  |
| --- | --- | --- | --- | --- |
|  | BSJs | Circular isoforms | BSJs | Circular isoforms |
| C666-1 | 15,812 | 19,280 | 4,842 | 3,758 |
| NPC43 | 22,090 | 29,648 | 5,368 | 4,236 |
| C15 | 8,401 | 8,350 | 3,145 | 2,247 |
| C17 | 16,131 | 22,422 | 4,505 | 3,396 |
| Xeno-32 | 17,270 | 17,888 | 4,966 | 3,763 |
| Xeno-666 | 19,809 | 21,412 | 5,118 | 3,976 |
| Xeno-1915 | 13,290 | 15,104 | 4,145 | 3,138 |
| Xeno-2117 | 13,465 | 16,101 | 4,332 | 3,278 |
| Xeno-99186 | 11,432 | 13,113 | 3,938 | 2,964 |
| NPC-M1 | 15,526 | 17,148 | 4,953 | 3,837 |
| NPC-M2 | 28,809 | 32,959 | 5,862 | 4,637 |
| NP69 | 14,708 | 18,161 | 5,160 | 4,162 |
| NP361 | 9,574 | 9,965 | 4,580 | 3,553 |
| NP460 | 16,998 | 18,611 | 5,681 | 4,562 |
| NP550 | 12,482 | 14,699 | 5,141 | 4,129 |

Taking all the NPC and NP samples together, 6,716 unique frequently expressed BSJs were identified. Looking up these BSJs from three circRNA databases, namely CIRCpedia v2 [41], CSCD [44] and MiOncoCirc v0.1 [18], we found that 5,786 of them (86.2%) were contained in at least one of these databases, while the remaining 930 (13.8%) were novel (Figure 5a). Due to our definition of frequently expressed BSJs, each of these novel BSJs was called in at least three, and usually more samples (Figure 5b), suggesting that these novel BSJs are frequently expressed in NP and NPC in a tissue- or cancer type-specific manner. We also checked the average number of supporting reads for each of these novel BSJs among the samples from which it was called, and found that around half of these novel BSJs had an average supporting read of 5 or more (Figure 5c), further supporting the reliability of these BSJ calls.

**Figure 5:**
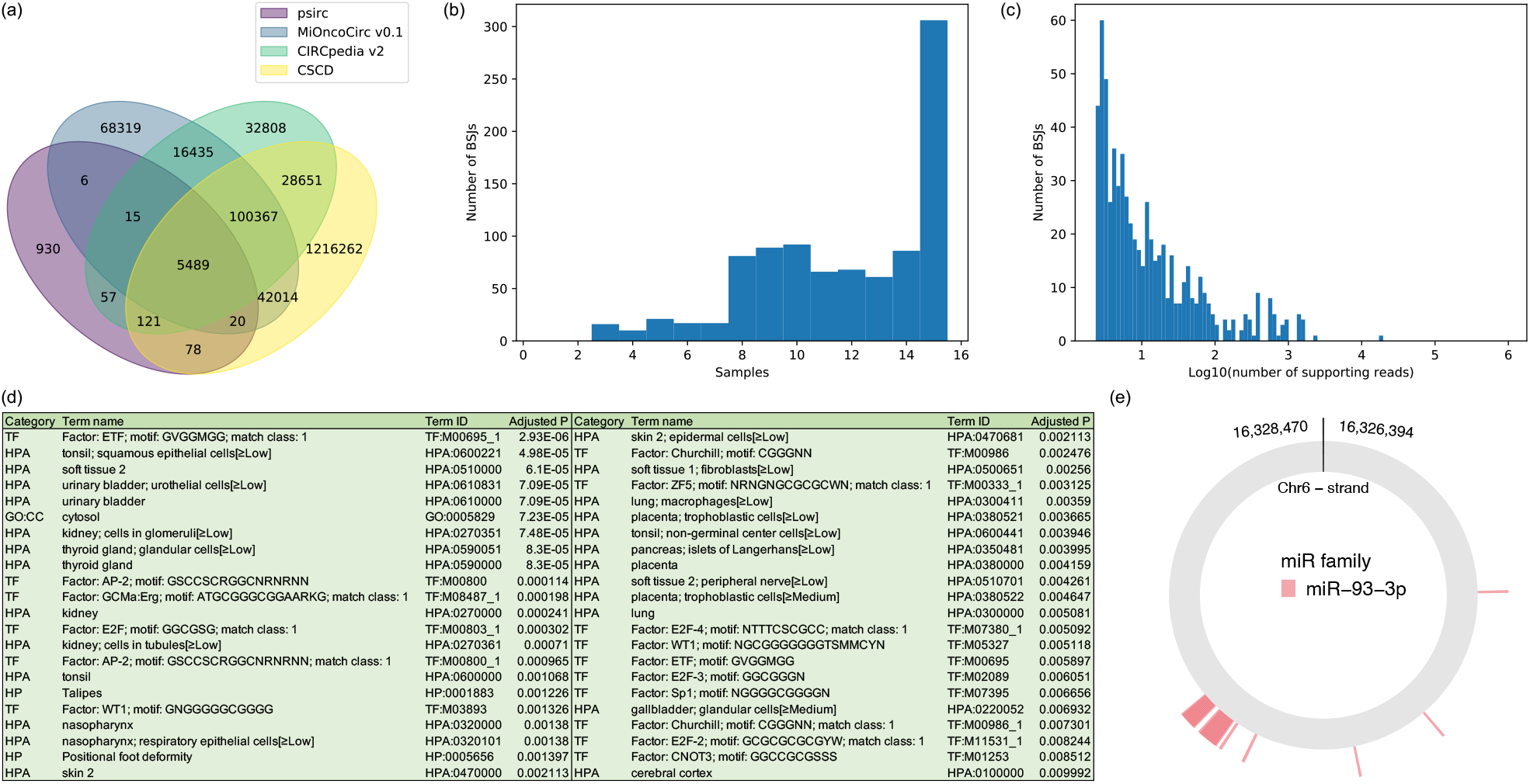
Analyses of the results produced by psirc from the NPC and NP samples. (a) A Venn diagram comparing the frequently expressed BSJs identified from the NPC and NP samples with those in three circRNA databases. (b-c) Histograms of the frequently expressed novel BSJs not contained by any of the three databases, in terms of the number of NPC and NP samples from which they were called (b) and their average supporting reads among the samples from which they were called (c). (d) Enriched functional terms (adjusted P *<* 0.01) of the down-regulated genes based on the full-length circRNA isoform analysis. (e) MREs on the differentially expressed *ATXN1* circRNA.

To see how the expression levels of linear and circular isoforms are related to each other, from each sample we extracted all pairs of linear and circular isoforms with identical sequences with at least one of them expressed. These pairs reveal a general positive correlation between the linear and circular expression levels, but some variations do exist (Figure S7). This is consistent with observations from previous studies based on specific examples or BSJs [11].

We then performed differential expression analysis, and obtained 155 BSJs and 319 full-length circRNA isoforms, coming from 129 and 271 genes, respectively, with relatively strong differential expression between the NPC and NP samples (Wilcoxon q-value ≤ 0.2, Supplementary Files 2, 4). Among these genes, 9 from the BSJ list and 57 from the full-length list were up-regulated in NPC, 8 of which were common. The numbers of down-regulated genes were larger, with 120 from the BSJ list and 214 from the full-length list, 86 of which were common. This trend of reduced circRNA expression in NPC is consistent with similar observations in prostate cancer and colorectal cancer [48, 49]. The differences between the BSJ and full-length lists show that some differentially expressed full-length circRNA isoforms would be missed if only the BSJs were quantified. Next, we performed a functional enrichment analysis of the differentially expressed genes on the full-length list using g:Profiler [50]. For the up-regulated genes, there were no significantly enriched functional terms. For the down-regulated genes, a fairly large number of functional terms were significantly enriched (Figure 5d), including Human Protein Atlas [51] terms “tonsil; squamous epithelial cells” (adjusted p=4.98E-5), “thyroid gland” (adjusted p=8.30E-5) and “nasopharynx” (adjusted p=1.38E-3).

Among the full-length circRNA isoforms with a differential expression p-value ≤ 0.05, we found 24 of them with at least 10 MREs of a single miRNA family (Table S4, Materials and Methods). Among them, a circRNA generated by back-splicing of an exon of the *ATXN1* gene was predicted to harbor 29 MREs of the miRNA family miR-93-3p (Figure 5e).

To further explore the significance of quantifying full-length circRNA isoforms, we compared the MREs of each frequently expressed circRNA isoform involving exon skipping with the corresponding isoforms derived from all annotated linear isoforms that contain the two exons defining the BSJ. We call the latter the “default isoforms” and the former the “alternative isoform”. In total, from 252 frequently expressed alternative isoforms, we derived 276 default isoforms. On average, each of these alternative isoforms harbors 45.6 fewer MREs than their default isoforms. In one extreme example, an alternative isoform harbors 10 fewer MREs from the same miRNA family than its default isoforms. Importantly, 31.2% of these alternative isoforms had an expression level over 100 times higher than the corresponding default isoforms. These results show that if full-length circRNA isoforms are not inferred and quantified but rather default isoforms are assumed based on BSJs and annotated linear isoforms alone, functional studies of circRNA could be seriously misinformed.

Finally, we experimentally verified some of the differentially expressed circRNAs between the NPC and NP groups. We started with a verification of the BSJs using RT-PCR (Figure 6a). The results are highly consistent with the TPM values of these BSJs determined by psirc based on the supporting reads (Figure 6b). For example, the two BSJs from *NETO2* were mostly expressed in the NP group but not the NPC group according to both RT-PCR and psirc. In contrast, the BSJs from *NTRK2* and the EBV-encoded *RPMS1* were mostly expressed in the NPC group but not the NP group according to both methods.

**Figure 6:**
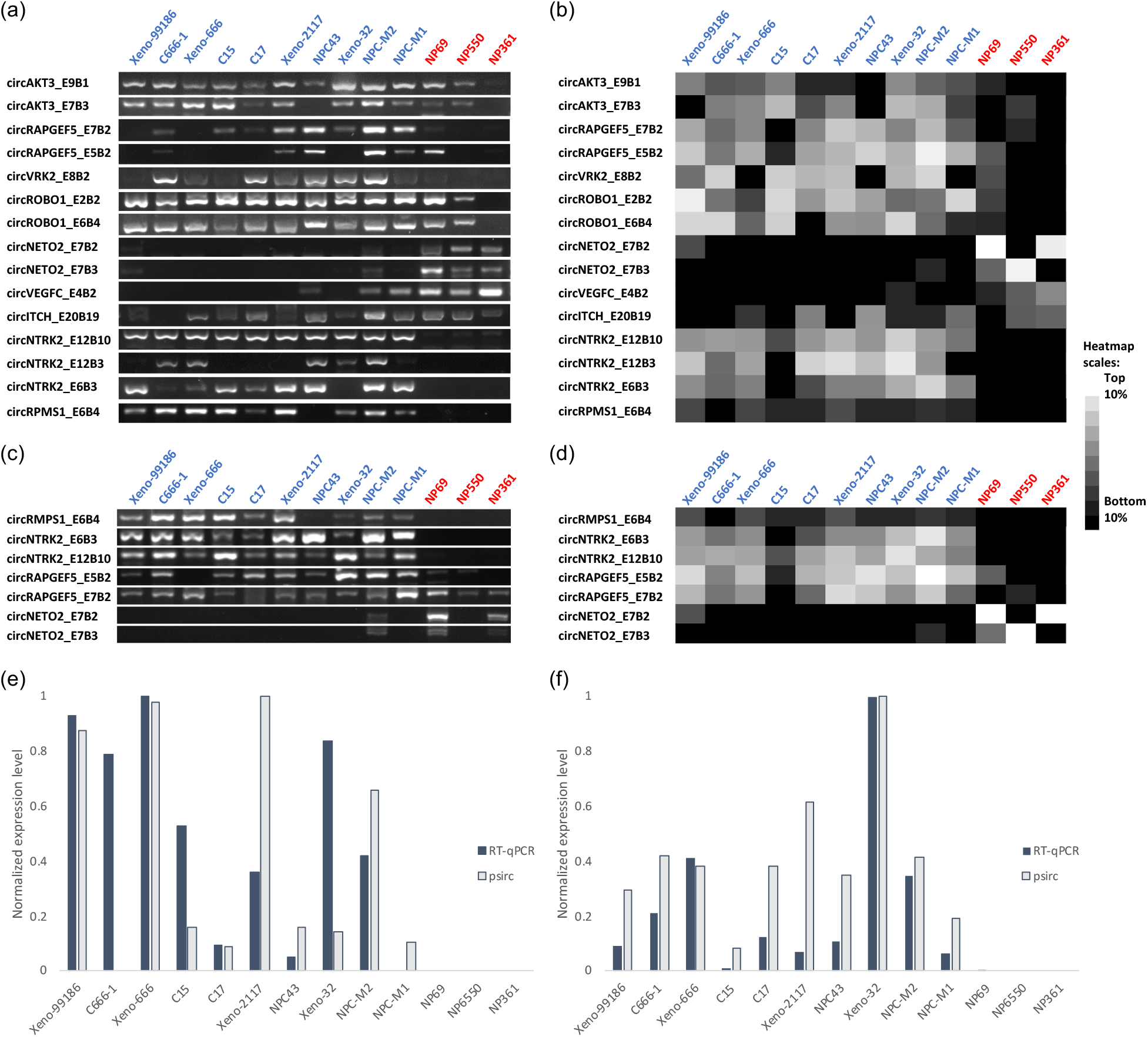
Experimental validations of computational results of psirc. (a) Validation of differentially expressed BSJs using RT-PCR. Each row corresponds to a BSJ and each column corresponds to a sample, with the NPC samples labeled in blue and the NP samples labeled in red. (b) Expression levels of the same BSJs determined by psirc. The expression value of each BSJ was computed by summing up the TPM values of all transcripts that involved this BSJ. (c) Validation of differentially expressed full-length transcript isoforms using RT-PCR. (d) TPM values of the same full-length isoforms inferred by psirc. (e-f) RT-qPCR results of two short full-length transcript isoforms, circRPMS1 E6B4 (e) and circNTRK2 E12B10 (f). In each of these two panels, normalization was performed by dividing each value by the largest value among all samples.

Next, we designed primers specifically for some differentially expressed full-length transcript isoforms. The RT-PCR results (Figure 6c) again show high consistency with the corresponding TPM levels deduced by psirc in distinguishing between the two sample groups (Figure 6d). To get a more quantitative evaluation of the deduced expression levels, we further performed RT-qPCR on two short full-length isoforms, selected according to the detection limit of RT-qPCR. The results (Figure 6e,f) show that the two isoforms had almost no expression in the NP group based on both the psirc and RT-qPCR results. When considering only the NPC samples, the two sets of results correlated positively (Pearson correlation = 0.41 for circRPMS1 E6B4 and 0.83 for circNTRK2 E12B10), with some differences between them likely caused by a combination of technical and biological reasons such as cell passages.

## 3 Discussion

In this study, we have developed psirc, the first complete pipeline that can identify both BSJs and full-length circRNA transcript isoforms of all lengths and quantify their expression levels. We have demonstrated the effectiveness and computational efficiency of psirc using RNA-seq data from human cell lines and fetal tissue samples. At the BSJ level, psirc achieved comparable sensitivity and precision as the best of CIRCexplorer2, CIRI2 and CircMarker while requiring substantially less running time and memory. At the full-length isoform level, psirc detected a lot more isoforms than CIRI-full and produced more accurate expression level quantification than Sailfish-cir, especially in terms of the relative expression levels of linear and circular transcripts.

The efficiency of psirc is due to its use of pseudo-alignment by kallisto, which avoids time-consuming full alignments of sequencing reads but is still able to accurately determine the transcript isoforms from which each sequencing read could come from by using the T-DBG. In contrast, the k-mer-based method of Sailfish-cir can assign reads to wrong transcript isoforms by ignoring the order of k-mers.

We have found that for some differentially expressed circRNA isoforms between NPC and NP, their expression levels were much higher than default isoforms assumed by “circularizing” annotated linear isoforms that contain the two exons defining the BSJ. The highly expressed alternative isoforms could have far fewer MREs than the default isoforms due to exon skipping. Being able to identify full-length circRNA isoforms and quantify their expression levels thus enable much more accurate study of potential circRNA functions such as their interactions with miRNAs and RNA binding proteins.

A recent study has shown that circRNAs can form 16-26bp duplex structures, which act as inhibitors of double-stranded RNA-activated protein kinase (PKR), and these circRNAs are degraded by RNase L for activating PKR during early innate immune response [13]. The structures of circRNAs and their corresponding functional mechanisms are still under investigation, but the full-length sequences of circRNAs likely play some roles in determining the possible structures.

By considering the read supports of individual forward-splicing and back-splicing junctions, psirc is able to infer circular transcript isoforms that contain an exon combination different from any annotated linear isoforms. However, psirc still relies on an input set of linear transcript isoforms to define exon boundaries. As a result, it cannot detect non-exonic circRNAs such as intron-exon circRNAs (elciRNAs) or circRNAs that involve alternative 5’ or 3’ splice sites. This limitation can be potentially overcome by first performing a linear transcript assembly on the RNA-seq data and augment the transcript annotation set with the novel transcripts identified, at the expense of extra computational resources.

## 4 Materials and Methods

### 4.1 Details of the psirc method

#### 4.1.1 Identification of BSJs

In the first step of psirc (Figure 1a), to detect junction-crossing reads, we construct a library of all possible BSJ sequences according to the transcripts defined in a gene annotation set. In the default setting of psirc, Gencode [52] (v29) is used. For each annotated linear transcript, we consider each pair of exons in turn to construct a sequence that contains the *x* nucleotides at the 3’ end of the 3’ exon in the pair followed by the *x* nucleotides at the 5’ end of the 5’ exon, and add the sequence to the library. The value of *x* is determined according to the size of k-mers (length-k sub-sequences) used in pseudo-alignment as a trade-off between the sensitivity and specificity of BSJ detection. In the default setting of psirc with a k-mer size of 31, *x* is set to 24 such that each of the two exons has at least 7bp of the k-mer pseudo-aligned to. All the BSJs with pseudo-aligned sequencing reads are then collected.

To gain further support for these BSJs, we construct a new sequence library as follows. For each exon pair that defines a detected potential BSJ, we take the corresponding full linear transcript sequence and shift *y* nucleotides at the 3’ end of it to the 5’ end, such that reads that support the BSJ can be pseudo-aligned normally without split-alignment. This rotated sequence is then added to the sequence library. The value of *y* is chosen to be the most common read length in the RNA-seq data set. All read pairs with one or both reads that supported BSJs based on the first sequence library are then pseudo-aligned to this second sequence library in paired-end mode. BSJs that are no longer supported by any reads after this step are removed from the list of potential BSJs.

Similarly, to detect the last-first exon read pairs, we construct a library containing all pairs of first and last exons of the same transcripts. In theory, as long as the two reads in a pair are pseudo-aligned to different exons of a transcript in an outward-facing manner, there is a potential BSJ. However, the exact location of the junction cannot be certain unless the two exons involved are the first and last ones of a transcript, in which case it must be the end of the last exon back-splicing to the beginning of the first exon, and therefore we only focus on this special case. Any such pairs with two reads in a pair pseudo-aligned in the outward-facing orientation would be considered a supported BSJ.

In all the above processes, pseudo-alignment is performed using kallisto, which determines the set of all sequences that could have produced the read. This is done by comparing k-mers in the read with the k-mers in the library sequences. This pseudo-alignment step in psirc is very fast because i) the total length of the library sequences is small as compared to the whole genome or transcriptome, ii) kallisto uses a hash table to efficiently map each k-mer to the sequences that contain it (called its “k-compatibility class”), and iii) kallisto uses a T-DBG to determine the k-mers that do not need to be checked when they belong to the same k-compatibility class and appear on the same non-branching path in the graph. Although the pseudo-alignment results do not contain full alignments between the reads and the library sequences at the per-nucleotide resolution, they do contain the aligned location(s) of each read, which is sufficient for our purpose of identifying BSJs. By default, we allow each read to be pseudo-aligned to at most 10 different locations. The version of kallisto used in psirc is a forked version that we modified from the main trunk, which can support multi-threading and produce a SAM file as output.

#### 4.1.2 Identification of potential full-length circular transcript isoforms

In the second step of psirc (Figure 1b), for each linear transcript isoform with a BSJ detected in the first step, we identify a set of potential full-length circular transcript isoforms as follows. Suppose the original linear transcript isoform in the annotation file involves exons *E*_1_, *E*_2_, …, *E*_*n*_, and in the first step of psirc, a BSJ was detected between exons *E*_*i*_ and *E*_*j*_, where 1 ≤ *i* ≤ *j* ≤ *n*. A directed graph is constructed with each exon *E*_*i*_, *E*_*i*+1_, …, *E*_*j*_ forming a node. For each pair of nodes *E*_*a*_ and *E*_*b*_ where *i* ≤ *a < b* ≤ *j*, if the forward splicing junction from *E*_*a*_ to *E*_*b*_ is supported by sequencing reads, an edge will be drawn from the former to the latter. In addition, edges are also added for exons that are adjacent in the original linear transcript isoform, i.e., from *E*_*a*_ to *E*_*a*+1_ for 1 ≤ *a < n*. A depth-first search is then performed to identify all non-cyclic paths from *E*_*i*_ to *E*_*j*_, and the nodes on each of these paths will form a potential full-length circular transcript isoform. Finally, if *i* = *j*, a potential full-length circular transcript isoform involving this exon alone will also be added to the list.

Practically, to avoid an excessive amount of computation spent on a single transcript, we did not produce the set of potential circular transcript isoforms for each linear transcript that had more than 20 read-supported forward splicing junctions between the two exons that defined the BSJ. We also omitted isoforms with at least five read-supported forward-splicing junctions involving non-adjacent exons. In our tests, only a very small number of BSJs satisfied these filtering criteria in the benchmark data sets and got excluded.

To ensure the reliability of identified isoforms involving non-adjacent exons (i.e. the “alternative isoforms”), those that have the exact same sequence as any identified non-alternative isoform are filtered out. Also, in the final output, isoforms with identical sequences coming from the same type (linear or circular), same genomic locus, and same strand are merged into one entry.

#### 4.1.3 Expression quantification of full-length transcript isoforms

In the third step of psirc (Figure 1c), the expression level of each linear and circular transcript isoform is estimated based on likelihood maximization. The overall workflow involves indexing, pseudo-alignment and quantification (Figure S8).

In the indexing stage, all the potential linear and circular transcript isoforms, including the ones originally in the annotation file of linear transcript isoforms and the ones identified in the second step of psirc, are used to construct a colored transcript de Bruijn graph (T-DBG). In the graph, each node corresponds to a k-mer and there is an edge from one node to another if the last k-1 nucleotides of the former are the same as the first k-1 nucleotides of the latter. Each transcript isoform is given a unique “color”, and each node is given a set of colors corresponding to the transcript isoforms that contain the k-mer. In this formulation, each linear transcript isoform corresponds to a path (possibly with repeated nodes and edges if some k-mers appear multiple times on the transcript sequence) and each circular transcript corresponds to a closed path. Since standard gene annotation files (such as those in GFF or GTF format) cannot handle circular transcript isoforms, we record the information of these isoforms directly in the T-DBG by adding edges caused by the BSJs. For example, suppose there is a circular transcript isoform that contains “ATC” at the beginning of its first involved exon and “GTC” at the end of its last involved exon. If *k* = 3, there will be edges added from “GTC” to “TCA”, from “TCA” to “CAT” and from “CAT” to “ATC”.

Nodes that are on the same linear path without any branching and share the same color are condensed into a single contig. After this condensation, the final form of the T-DBG is a graph of contigs each associated with a set of colors, which is also called the equivalent class (EC) of the contig.

In the pseudo-alignment stage, all sequencing reads are pseudo-aligned to the transcriptome defined by the T-DBG containing both linear and circular transcript isoforms. If the sequence of a read can be found on multiple transcript isoforms, it would be pseudo-aligned to all of them. In particular, if a linear transcript isoform and a circular transcript isoform share exactly the same set of exons, sequencing reads produced from them that do not cross the BSJ would be pseudo-aligned to both. Only the reads that cross the BSJs can be uniquely pseudo-aligned to the circular transcript isoforms. In the original implementation of kallisto, the relative orientation and distance between paired reads in paired-end sequencing data are not considered. We extended it by using the orientation and insert size information to validate the pseudo-alignments of the individual reads and to check whether the sequenced RNA fragment comes from a circular transcript. The final output of this stage is a list of equivalent classes and the corresponding number of sequencing reads pseudo-aligned to each of them.

In the quantification stage, the expression level of each transcript isoform, defined as the number of transcript copies, is quantified by maximizing the data likelihood according to a probabilistic model [47] under the assumption that reads are correctly pseudo-aligned. Two types of biases need to be corrected, both by adjusting the effective length of transcripts. The first type of bias is caused by a drop of sequencing depth at the ends of linear transcripts, since fragments produced near the end of a transcript could be too short to be sequenced by the experimental procedure. This type of bias does not apply to circular transcripts, assuming that circular transcripts can be cut anywhere with equal chance. To correct for this bias, the effective length of each linear transcript is defined as its actual length minus the average fragment length (i.e., half the fragment length from each end). The second type of bias is caused by the GC content of transcripts and the corresponding non-uniform sequence amplification. To correct for this bias, the occurrence counts of all 6-mers on the sequencing reads are compared with their expected occurrences based on the estimated abundance of each transcript isoform, as proposed previously [47].

### 4.2 Data sets for testing psirc’s performance

We obtained three RNA-seq data sets for testing psirc’s performance and compared it with other circRNA detection methods (Table S1). The first data set contained rRNA-depleted RNA-seq data of human fetal samples [29] (GEO accession: GSE64283). Among the 35 samples in this data set, 11 of them had RT-qPCR measurements of the expression of BSJs from 8 genes, with 79 measurements in total. We used only these 11 samples in our analyses, and downloaded the corresponding RNA-seq data from Sequence Read Archive [53]. The second and third sets of data contained rRNA-depleted RNA-seq and RNase R-treated RNA-seq data of HeLa and Hs68 cells [54, 55]. Replicate samples were combined by pooling the data directly.

### 4.3 Other circRNA calling methods compared

We compared psirc with a number of existing methods, including CIRCexplorer2 [34] (v2.3.8), CircMarker [27] (git commit 06aa680a), CIRI2 [23] (v2.0.6), CIRI-full [36] (v2.0, with CIRI AS v1.2 and CIRI-vis v1.4), CIRIquant [35] (v1.1) and Sailfish-cir [26] (v0.11, with sailfish v0.9.2). All these methods were run with their default parameter values.

We categorized these methods based on their abilities to identify BSJs, infer full-length transcript isoforms, and quantity the expression of them (Table S2). For methods that infer and quantify full-length transcript isoforms, BSJ read counts were computed as 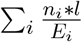, where *n*_*i*_ is the read count of transcript *i, l* is the read length, *E*_*i*_ is the effective length of transcript *i*, and the summation is over all transcripts that involve the BSJ. In our analyses we considered both raw read counts and normalized read counts, defined as raw read counts per million aligned reads.

### 4.4 Testing psirc’s ability to identify BSJs

We applied CIRCexplorer2, CIRI2, CircMarker and psirc to identify BSJs from the three data sets as explained in Results. For CIRCexplorer2, we failed to run it in its default setting on the sample Hs68 C1. Therefore, we instead downloaded the CIRCexplorer2 results of HeLa and Hs68 from the supplementary materials (“Presentation2.zip”) of Hansen (2018) [38] and used these results in the comparisons directly.

For the analyses involving the HeLa and Hs68 data, for each BSJ identified, we classified it into either Enriched, Unaffected, Depleted or Abolished, according to the numbers of reads that support this junction in the rRNA-depleted data and RNase R-treated data. Following previous studies [37–39], unnormalized read counts were used to define these classes. For each of the two samples, since the total number of read pairs after pooling the data form replicates is highly consistent with or without the RNase R treatment (Table S1), using unnormalized read counts should not create any strong systematic bias. For a particular BSJ, suppose the number of supporting reads in the rRNA-depleted data is *d*, the number of supporting reads in the RNase R-treated data is *t*, and *α* is an enrichment factor [37, 38], the definitions of the four classes are as follows:

- Enriched: *t* ≥ *d* × *α*
- Unaaffected: *d* × *α* > *t* ≥ *d*
- Depleted: *d* > *t* > 0
- Abolished: *t* = 0

We used the enrichment factor *α* = 1.5 for HeLa and *α* = 5 for Hs68. The value for Hs68 was taken directly from previous studies [37, 38], while the value for HeLa was the total number of identified BSJ reads in the RNase R-treated samples divided by that in the untreated samples.

While the Enriched cases were likely true positives and the Abolished cases were likely false positives, the Unaffected and Depleted cases were more ambiguous. We therefore considered four different measures of precision in order to provide a comprehensive evaluation of the performance of the four methods. In all four definitions, precision was defined as TP/(TP+FP), where TP stands for true positives and FP stands for false positives. The four precision measures differed by their definitions of TP and FP:

- Precision 1: TP involved the Enriched cases only, FP involved the Abolished cases only
- Precision 2: TP involved the Enriched and Unaffected cases, FP involved the Abolished cases only
- Precision 3: TP involved the Enriched cases only, FP involved the Depleted and Abolished cases
- Precision 4: TP involved the Enriched and Unaffected cases, FP involved the Depleted and Abolished cases

We quantified the computational costs by elapsed time, CPU time, and RAM usage. Elapsed time was defined as the duration of the physical running time, from the time that a method started to the time that it completed. CPU time was the total amount of time used by the CPU on all the threads of the method. These two time measurements differed mainly in two aspects, namely 1) elapsed time could be shortened by multi-threading, but CPU time was the total of all threads, and 2) elapsed time included all the overheads such as disk I/O, but CPU time just included the time spent on CPU cycles. RAM usage was defined as the peak memory usage during the whole execution process.

All the methods were run on a machine with 64 Intel Xeon E5-4610 v2 cores@2.30GHz with 520GB of RAM. At any time only one method was run on one sample without any other user processes running in the background.

### 4.5 Comparing with RT-qPCR measurements

Expression values deduced from RNA-seq data were log_2_-transformed after addition of a small constant (*c*) to handle the zero-expression cases. The value of *c* was set to 1 and 0.01 for raw and normalized BSJ read counts, respectively. These values were chosen because they were close to the smallest non-zero values observed.

### 4.6 Generation of simulated RNA-seq data

Sequencing reads of linear and circular transcripts were generated based on the RefSeq gene annotation of the human genome reference GRCh38.p12. We first extracted from the annotation all 23,851 genes with at least one transcript isoform containing a CDS feature. We generated two sets of data based on these genes, namely an easier set with only one linear transcript isoform and one circular transcript isoform per gene, and a second set with more complex gene structures. In both cases, transcript copy numbers were drawn independently for each transcript isoform from the uniform distribution *U* (0, 1000). 9,000,000 fragments were generated for each group of genes (see below) according to these transcript copy numbers with a normal length distribution *N* (250, 25), and these fragments were then used to generate paired-end sequencing reads of 100bp long using Polyester-LC, which was an extension of Polyester [56] we made to generate sequencing reads of both linear and circular transcripts. The only difference between Polyester-LC and Polyester is that for circular transcripts, Polyester-LC can determine the starting and ending points of each fragment uniformly without the end effect of linear transcripts. In order to quantify the complexity of each gene, we introduced a metric called Transcript Overlap Ratio (TOR), defined as the number of linear transcript isoform pairs with overlapping regions divided by the total number of linear transcript isoforms.

For the first set of data, we first sorted all genes with at least 10 transcript isoforms containing CDS features by their TOR values. Starting from the genes with the lowest TOR values, we randomly selected one linear transcript isoform of length (considering only the exons) at least 900bp from each gene (if there is any) until we have collected 500 linear transcript isoforms in total. Based on these linear transcript isoforms, we then generated a linear transcript isoform and a circular transcript isoform of the same length with a certain overlap ratio between them, where the overlap ratio ranged from 0% to 100%, forming 11 groups of data. Specifically, the synthetic linear transcript isoform occupies the beginning of the real linear transcript isoform while the synthetic circular transcript isoform occupies the end, with their lengths determined according to the target overlap ratio (Figure S6). For example, when the overlap ratio was 0%, the synthetic linear transcript isoform was equal to the first half of the real linear transcript isoform and the synthetic circular transcript isoform was equal to the second half. As a result, this data set contained 11 groups each with 1,000 synthetic transcript isoforms including 500 linear ones and 500 circular ones. Sequencing reads were then generated from these 11 groups of transcript isoforms as explained above.

For the second set of data, we collected the first 100 genes on the TOR-sorted gene list each with at least 20 linear transcript isoforms. We then randomly sampled a certain number of transcript isoforms from each gene, where the number ranged from 1 to 10, forming 10 groups. For each linear transcript isoform sampled, we also synthesized a corresponding circular transcript isoform with the same sequence as the linear counterpart. As a result, we had 20-200 transcript isoforms per group, and sequencing reads were generated from them according to the procedure described above. We repeated the whole process 10 times with 10 sets of random transcript isoforms sampled from the 100 genes.

### 4.7 Production and processing of RNA-seq data from the NPC and NP samples

Four immortalized normal nasopharyngeal epithelial cell lines (NP69, NP361, NP460 and NPC550), 2 EBV-positive NPC cell lines (C666-1 and NPC43), 7 patient-derived xenografts (Xeno-666, Xeno-2117, Xeno-1915, Xeno-99186, C15, C17 and Xeno-32) [57–62] and 2 patient tumor specimens (NPC-M1 and NPC-M2) were used in this study. NPC tumor specimens were from patients admitted to Prince of Wales Hospital, The Chinese University of Hong Kong. Patient consents were obtained according to institutional clinical research approval (IRB) at The Chinese University of Hong Kong, Hong Kong SAR. Total RNA extracted from the NP and NPC samples were subjected to rRNA depletion by Ribo-Zero kit (Illumina), followed by TruSeq stranded total RNA library construction and sequencing on the Illumina Hi-seq2000 system according to the manufacturer’s protocols [62].

### 4.8 Information contained in the output files of the NPC data analysis

The results of the NPC data analysis are summarized in four files, namely a file of all detected BSJs, an annotation file of all candidate full-length linear and circular transcript isoforms, a file of the corresponding sequences of these full-length isoforms, and a file that provides the quantification results of these isoforms.

In the first file, provided in a tab-delimited format, each line lists a BSJ indexed by the two genomic positions right next to the junction in the two respective exons. All candidate full-length transcript isoforms that contain the BSJ are also listed, which serve as a mapping between this file and the second file. The number of reads that support the junction in each sample is listed next. The mean supporting read count in each of the two sample groups is also shown, together with the log fold change of these mean values and the corresponding differential count p-value based on Wilcoxon rank-sum test and its q-value. To focus on the most interesting cases, only junctions with a non-zero read count in at least 70% NPC or NP samples are included.

In the second file, provided in the GTF format, each line lists an exon and the corresponding genes and transcript isoforms that contain it. The ID of each circular transcript isoform ends with the exon connection information in the form of “ _E*w*B*x{ −y*_*i*_A*z*_*i*_*}*\*”, where *w* and *x* indicate the 3’ and 5’ exons that define the BSJ, and each pair of *y*_*i*_ and *z*_*i*_ defines a junction that involve non-adjacent exons, and there can be zero, one or more of them. For example, “HP1BP3_E9B1_2A4” means that this circular isoform involves a BSJ from exon 9 of the HP1BP3 gene back to exon 1, and exon 2 is connected to exon 4 directly skipping exon 3, followed by exons 4 to 9.

In the third file, provided in the FASTA format, the actual sequences of the full-length transcript isoforms defined in the second file are listed.

In the fourth file, provided in a tab-delimited format, each line lists a candidate full-length transcript isoform and its expression level in each of the samples in TPM (transcripts per million reads aligned) values. The mean expression in each of the two groups, log fold change, differential expression p-value and q-value are also provided. Again, only transcript isoforms with a non-zero read count in at least 70% of NPC or NP samples are included.

### 4.9 Comparing the frequently expressed BSJs identified from the NPC and NP samples with those in circRNA databases

The frequently expressed cellular BSJs identified from the NPC and NP samples were combined and de-duplicated, resulting in a set of 6,723 unique BSJs. These BSJs were compared with those from three circRNA databases. For CIRCpedia v2, a total of 183,943 unique BSJs from all human cell lines were downloaded from https://www.picb.ac.cn/rnomics/circpedia/. For CSCD (accessed on June 9, 2020), the union of all common, normal and cancer-specific BSJs was taken, all downloaded from http://gb.whu.edu.cn/cscd/, resulting in a set of 1,393,002 unique BSJs. For MiOncoCirc v0.1, BSJs from all cancer samples were downloaded from https://mioncocirc.github.io/download/, with a total of 232,665 unique BSJs. All genomic positions in the four sets were based on the human reference genome GRCh38.

### 4.10 Functional enrichment analysis

We used g:Profiler to perform functional enrichment analysis of the differentially expressed circRNAs obtained from the full-length quantification results of psirc. We ran g:Profiler using its default setting, which included the following functional categories: Gene Ontology sub-ontologies (molecular function, cellular component, and biological process), biological pathways (KEGG, Reactome, and WikiPathways), regulatory motifs in DNA (TRANSFAC and miRTarBase), protein databases (Human Protein Atlas and CORUM), and Human phenotype ontology (HP).

### 4.11 Prediction of MREs

Information about high-confidence human and EBV miRNAs and their families was downloaded from miRBase (release 22) [63]. In total, 897 human miRNAs from 736 families and 44 EBV miRNAs from 44 families were involved. Different miRNAs in the same family share the same seed. One EBV miRNA (ebv-miR-BART4-3p) was found to have the same seed as a human miRNA (hsa-miR-499a-3p). Therefore, altogether 736+44-1 = 779 miRNA families were considered. MREs on potential circRNA sequences were identified using TargetScan (v7.0) [64] with default parameter values. All bases on the sequences were not masked, which allowed MREs to appear anywhere on the sequences. To allow for detection of MREs that overlap the BSJs, 10 bases from the 5’ end of each sequence were copied and pasted to the 3’ end. Redundant MREs that appeared completely within the 10 bases were removed to avoid double counting.

### 4.12 Experimental validations

The candidate BSJs and full-length isoforms of selected differentially expressed cellular and viral circRNAs of NPC and NP samples identified by psirc were subjected to conventional RT-PCR analysis and RT-qPCR analysis. The predicted BSJs of cancer gene-derived circRNAs either predominantly over-expressed (e.g. circAKT3, circVRK2, circNTRK2) or down-regulated (circNETO2, circVEGFC) in multiple NPC tumor samples were selected for validation by conventional RT-PCR with primers flanking the junction. For validation of full-length isoforms predicted by psirc, conventional RT-PCR analysis with inverse primer pair in the same exon for selected over-expressed EBV-encoded (circRMPS1) and cellular (cirNTRK2, circRAGEF5) circRNAs, as well as down-regulated circRNAs (circNETO2), was performed in a panel of NPC and NP samples. For the RT-PCR experiments, total RNA of the samples was extracted with TRIzol Reagent (Invitrogen, USA) and then subjected to cDNA synthesis with SuperScriptTM III Reverse Transcriptase (Invitrogen) and PCR amplification with KAPA2G Fast HotStart PCR Kit (Roche, USA) according to the manufacturer’s protocol. PCR products were run on a 2% agarose gel. The primers involved in the validations of BSJs and full-length transcripts are listed in Table S5 and Table S6, respectively.

Two relatively small-sized circRNAs (circRPMS1-E6B4, 399 bp; circNTRK2-E12B10, 237 bp) were further validated by RT-qPCR analysis. cDNA samples were subjected to quantitative PCR analysis using the TaqMan Universal PCR Master Mix (Applied Biosystems) on a LightCycler 480 Instrument II as described [65]. The primers involved are listed in Table S7. The RT-qPCR results were analyzed by delta-delta Ct method and normalized by GAPDH expression.

## 5 Data Availability

The published RNA-seq data from previous studies were obtained from the NCBI Sequence Read Archive (SRA) as listed in Table S1. The RNA-seq data for the NPC and NP samples have been submitted to Gene Expression Omnibus [66], with the accession number to be assigned.

## 6 Code Availability

The source code of psirc is available at the website https://github.com/Christina-hshi/psirc.git.

## 7 Acknowledgement

This project was supported by the Hong Kong Research Grants Council Theme-based Research Scheme T12-401/13-R. YYL was supported by Fundamental Research Grant Scheme (FRGS/1/2017/SKK08/UM/02/11). KWL and CMT were supported by the Hong Kong Research Grants Council Area of Excellence AoE/M-401/20, Collaborative Research Fund C4001-18GF and General Research Fund 14113620. KYY was supported by Hong Kong Research Grants Council Collaborative Research Funds C4045-18WF, C4054-16G, C4057-18EF, and C7044-19GF and General Research Funds 14107420, 14170217 and 14203119, the Hong Kong Epigenomics Project (EpiHK), and the Chinese University of Hong Kong Young Researcher Award and Outstanding Fellowship.

